# Distinct localization patterns of actin microfilaments during early cell plate formation in plants through deep learning-based image restoration

**DOI:** 10.1101/2025.01.28.635247

**Authors:** Suzuka Kikuchi, Takumi Kotaka, Yuga Hanaki, Minako Ueda, Takumi Higaki

**Affiliations:** Graduate School of Sciences and Technology for Innovation, Yamaguchi University, Yamaguchi, Japan; Faculty of Science, Kumamoto University, Kumamoto, Japan; Graduate School of Life Sciences, Tohoku University, Sendai, Japan; Graduate School of Science and Technology, Kumamoto University, Kumamoto, Japan; International Research Organization for Advanced Science and Technology, Kumamoto University, Kumamoto, Japan; International Research Center for Agricultural and Environmental Biology, Kumamoto University, Kumamoto, Japan

**Keywords:** tobacco BY-2 cells, cytokinesis, phragmoplast, microtubules, actin microfilaments, live-cell imaging, image restoration, deep learning

## Abstract

Phragmoplasts are plant-specific intracellular structures composed of microtubules, actin microfilaments (AFs), membranes, and associated proteins. Importantly, they are involved in the formation and expansion of cell plates that partition daughter cells during cell division. While previous studies have revealed the important role of cytoskeletal dynamics in the proper functioning of the phragmoplast, the localization and role of AFs in the initial phase of cell plate formation remain controversial. Here, we used deep learning-based image restoration to achieve high-resolution 4D imaging with minimal laser-induced damage, enabling us to investigate the dynamics of AFs during the initial phase of cell plate formation in transgenic tobacco BY-2 cells labeled with Lifeact-RFP or RFP-ABD2 (actin binding domain 2). This computational approach overcame the limitation of conventional imaging, namely laser-induced photobleaching and phototoxicity. The restored images indicated that RFP-ABD2 labeled AFs were predominantly localized near the daughter nucleus, whereas Lifeact-RFP labeled AFs were found not only near the daughter nucleus but also around the initial cell plate. These findings, validated by imaging with a long exposure time, highlight distinct localization patterns between the two AF probes and suggest that Lifeact-RFP labeled AFs play a role in initiating cell plate formation.

## Introduction

The phragmoplast is a plant-specific intracellular structure that forms a cell plate to partition daughter cells during cytokinesis (Rasmussen et al., 2013; Smertenko et al., 2017). The backbone of the phragmoplast is made up of microtubules (MTs), which are involved in the assembly of the cell plate, together with actin microfilaments (AFs), membranes, and associated proteins (Müller and Jürgens, 2016; Smertenko 2018). Proper formation and expansion of the phragmoplast is crucial for the completion of plant cytokinesis (Livanos and Müller, 2019).

The formation and expansion of the phragmoplast are regulated not only by MTs but also by the dynamics of AFs. MTs in the phragmoplast serve as essential tracks for the transport of vesicles to the division site, facilitating the construction of the cell plate during plant cytokinesis (Jürgens, 2005). Previous studies on fixed or living cells have reported that AFs exhibit MT-related dynamics and contribute to cell division by regulating cell plate expansion in plants. AFs specifically labeled with fluorescent phalloidin and/or monoclonal antibodies have been observed in fixed cells, including endosperm cells of *Haemanthus katherinae* (Schmit and Lambert, 1987), root cells of *Allium cepa* (Clayton and Lloyd, 1985), suspension culture cells of *Daucus carota* (Traas et al., 1987), suspension culture cells of *Medicago sativa* (Seagull et al., 1987), tobacco BY-2 cells (Kakimoto and Shibaoka, 1987; Hasezawa et al., 1991), and suspension culture cells of *Arabidopsis thaliana* (Li et al., 2010). Moreover, AFs labeled through the microinjection of fluorescent phalloidin or expression of probes labeled with fluorescent proteins have been observed in living cells, including endosperm cells of *H. katherinae* (Schmit and Lambert, 1990), endosperm cells of *Clivia nobilis* (Endlé et al., 1998), stamen hair cells of *Tradescantia virginiana* (Cleary et al., 1992), and tobacco BY-2 cells (Sano et al. 2005; Yu et al. 2006, Higaki et al. 2008, Maeda et al., 2020). Interestingly, AFs located between the cortical division zone (CDZ) and the phragmoplast guide the edges of cell plates to the CDZ, playing a key role in the formation of the cell plate and selection of the division plane (Kojo et al., 2014; Arima et al., 2018). Thus, AFs regulate the centrifugal growth of the phragmoplast, playing an important role in cytokinesis.

The contribution of phragmoplast AFs to the initial phase of cell plate formation has been increasingly elucidated in recent years, although to a considerably lesser extent than their roles in the later stages of cytokinesis. We previously reported that pharmacological disruption of AFs by actin polymerization inhibitor bistheonellide A leads to malformation of the emerging cell plate in the initial phase of cytokinesis (Higaki et al., 2008). We also found that disruption of AFs by the action of actin polymerization inhibitor latrunculin B (LatB) inhibits the contraction of phragmoplast MTs during the initial phase of cytokinesis, affects the timing of accumulation of NACK1 kinesin, which plays an important role in cell plate expansion, and delays the accumulation of the cell plate membrane (Maeda et al., 2020; Maeda and Higaki, 2021). These results suggest that AFs contribute to the proper formation of the cell plate via the contraction of MTs in the initial phragmoplast. Moreover, these studies suggest that AFs are accumulated in the midplane of the mitotic apparatus during the initial phase of cell plate formation, in other words, near the emerging cell plate. However, this has not been clearly verified in previous studies using a probe of AFs, namely the actin binding domain 2 (ABD2) of AtFIM1 (Sano et al. 2005; Higaki et al., 2008; Maeda et al., 2020), likely because of the limitations of the imaging technique or the fluorescent probe used to label the AFs. Therefore, there is a need to further observe the localization of phragmoplast AFs in the initial phase of cell plate formation.

Precisely capturing the dynamics of phragmoplast AFs in the initial phase of cell plate formation remains challenging. In tobacco BY-2 cells, the progression from phragmoplast formation to expansion occurs within a few minutes (Maeda et al., 2020), necessitating live-cell imaging with high spatiotemporal resolution. However, in confocal laser microscopy, increasing spatiotemporal resolution by increasing the exposure time or shortening the interval of the acquisition time can lead to negative effects such as significant degradation of fluorescent proteins and cell cycle arrest owing to excessive laser irradiation (Supplementary Figure 1). Previous studies have also reported the negative effects of laser irradiation on fluorescent probes, such as photobleaching and phototoxicity, which are related to the exposure time and laser power (Magidson and Khodjakov, 2013; Skylaki et al., 2016; Ojha and Ojha, 2021). Although minimizing the exposure time and/or laser power reduces the negative effects of light irradiation on the sample, it also reduces the signal-to-noise ratio and degrades spatiotemporal resolution. Therefore, addressing the trade-off between minimizing sample damage and enhancing image quality is critical for detailed investigations of AF dynamics using live-cell imaging.

Here, we explored the possibility of solving this problem computationally. Recently, we demonstrated the effectiveness of deep learning in microscopy image transformation in cell and cytoskeleton segmentation (Kikukawa et al., 2021; 2023; Horiuchi et al., 2024) and virtual staining (Ichita et al., 2024). Building on these advancements, we applied a deep learning-based image restoration technique to enhance the quality of images acquired with short exposure times, enabling live-cell imaging with high spatiotemporal resolution while minimizing sample damage, such as photobleaching and phototoxicity. Using this strategy, we aimed to tackle the challenging task of investigating AF localization during the early stages of cytokinesis.

In this study, we performed three-dimensional time-lapse (4D) imaging of cytokinetic tobacco BY-2 cells using different AF probes, namely Lifeact-RFP and RFP-ABD2, and subsequently, analyzed the data using a deep learning-based image restoration technique. Deep learning-based image restoration enabled us to identify key spatiotemporal regions of interest from comprehensive datasets acquired with high spatiotemporal resolution and reduced cellular damage, namely photobleaching and phototoxicity. We then validated the AF localization patterns indicated by the image restoration results through targeted, long-exposure imaging without the use of deep learning. This direct observation confirmed that the localization of phragmoplast AFs varied depending on the type of fluorescent probe used. Specifically, we found that AFs labeled with Lifeact-RFP, rather than RFP-ABD2, were prominently localized on the midplane of the division apparatus in the initial phase of cell plate formation. These findings support a possible contribution of phragmoplast AFs to the initial phase of cell plate formation.

## Materials and Methods

### Tobacco BY-2 cell culture

Tobacco BY-2 cells (*Nicotiana tabacum* L. cv. Bright Yellow 2) were fluorescently labeled with YFP–β-tubulin (YFP-TUB6), histone H2B–dCFP, and Lifeact–mCherry (Lifeact–RFP) (BY- YTHCLR, Yasuhara and Kitamoto 2014) or YFP–β-tubulin (YFP-TUB6) and red fluorescent protein (tdTomato)-actin-binding domain of the AtFIM1 fusion protein (RFP-ABD2) (BY- YTRF, Kojo et al., 2014) to detect MTs and AFs. Transgenic tobacco BY-2 cells were cultured in modified Linsmaier and Skoog (LS) medium (Nagata et al., 1992). The medium contained the following components: 4.6 g/L Murashige and Skoog plant salt mixture (392–00591, Fujifilm Wako, Osaka, Japan), 30 g/L sucrose (193–00025, Fujifilm Wako), 0.2 g/L potassium dihydrogen phosphate (169–04245, Fujifilm Wako), 0.1 g/L myo-inositol (I5125, Sigma, St. Louis, MO, USA), 1.0 mg/L thiamin hydrochloride (205–00855, Fujifilm Wako), and 0.2 mg/L 2.4- dichlorophenoxyacetic acid (040–18532, Fujifilm Wako). The pH of the medium was adjusted to 5.8 with KOH. The medium was dispensed into 300 mL flasks (95 mL medium/flask). Each flask was capped with aluminum foil and autoclaved at 120 °C for 20 min. Appropriate volumes of BY- 2 cell suspension were dispensed into the sterilized medium (1 mL of BY-YTHCLR and 5.5 mL of BY-YTRF) and agitated in a rotary shaker at 130 rpm and 27 °C in the dark. The cell suspension was removed from the flask on day 7 of incubation and transferred to a new flask for subculturing.

### Microscopic observation of tobacco BY-2 cells

One day after passaging, transgenic tobacco BY-2 cells were transferred into φ35-mm Petri dishes with φ14-mm coverslip windows at the bottom (Matsunami, Osaka, Japan). The dishes were placed onto the inverted platform of a fluorescence microscope (IX-70; Olympus, Tokyo, Japan) equipped with a CSU-X1 scanning head (Yokogawa, Tokyo, Japan) and a scientific CMOS camera (Prime 95 B; Teledyne Photometrics, Tucson, AZ, USA). YFP was excited with a 488 nm laser and its fluorescence was detected with a 510–550 nm band-pass filter. In contrast, RFP was excited with a 555 nm laser and its fluorescence was detected with a 617–673 nm band-pass filter. The output power of each laser was set to 5% for YFP and 10% for RFP. Images were taken with a silicon-immersion objective lens (60×). The exposure time was adjusted according to the purpose of image acquisition and the fluorescence intensity of transgenic tobacco BY-2 cells.

### Microscopic observation of *A. thaliana* zygotes

Zygote imaging was performed using a marker of MTs (EC1p::Clover-TUA6, Horiuchi et al., 2024) and a laser-scanning inverted microscope (AX; Nikon) equipped with a pulse laser (InSight X3 Dual option; Spectra-Physics), as previously described (Kurihara et al., 2017). Fluorescence signals were detected using external non-descanned GaAsP PMT detectors, with a 525/50 nm band-pass filter, 40× water-immersion objective lens (CFI Apo LWD WI; Nikon), and Immersol W 2010 (Zeiss) immersion medium. Low-, medium-, and high-quality images of the same zygote were acquired by increasing the set laser power 2, 3, and 6 times, respectively.

### Acquisition of training image sets of tobacco BY-2 cells and model training

Training image sets were obtained by acquiring images of the same BY-YTHCLR cells at long-and short-exposure times. All images were cropped using ImageJ to create a region of interest (ROI) of 733 **×** 974 pixels. The pre-processed training images were used to train the image restoration model using image analysis software Aivia (DRVision, Bellevue, WA, USA) (Kikukawa et al., 2021; 2023). “2D Image Restoration” of Aivia was applied to train the model. For image training, a long-exposure image (Example) and a short-exposure image (Raw Data) of the same cell were used as a set. As a result, the model was trained on sets of “2000 and 20 ms images,” “2000 and 50 ms images,” “2000 and 100 ms images,” “2000 and 200 ms images,” and “2000 and 200 ms images.” The effect of the number of training image sets (N=2, 6, 12, 25, 50, and 100) on image restoration accuracy was examined.

### Acquisition of training image sets of *A. thaliana* zygotes and model training

Training image sets were obtained by acquiring images of the same *A. thaliana* zygotes with MTs fluorescently labeled with Clover-TUA6 at high, medium, and low laser powers (see Materials and Methods “Microscopic observation of *A. thaliana* zygotes” for the respective laser power). The training images were used to train the image restoration model using image analysis software Aivia (DRVision). “2D Image Restoration” of Aivia was applied to train the model. For image training, high-quality images (Example) and medium-or low-quality images (Raw Data) of the same zygote were used as sets, and the model was trained for each combination of “high laser power and medium laser power” and “high laser power and low laser power.” The number of training image sets was tested with N=25.

### Quantitative evaluation of image restoration accuracy

Quantitative evaluation of image restoration accuracy was performed by measuring the signal co-localization and correlation between the long-exposure image and the restored image from the short-exposure image. Specifically, Spearman’s rank correlation coefficients for “long-exposure images (2000 ms)” and “images restored from short-exposure images (20, 50, 100, or 200 ms)” were examined using ImageJ plugin EzColocalization (Stauffer et al., 2018).

### Three-dimensional (3D) visualization of time-lapse images

3D reconstruction was performed on Z-stack images at selected time points using Amira (Thermo Fisher Scientific, Pittsburgh, PA, USA). Z-stack images used for 3D reconstruction were processed by image restoration for each optical section prior to analysis in Amira. 3D images were visualized using the volume rendering function. GFP and RFP channels were displayed in pseudo color.

## Results

### The accuracy of image restoration by the model improves in proportion to the quality and number of training images

To investigate the localization of AFs in the initial phragmoplast under conditions of reduced photobleaching and phototoxicity, we decided to train the deep learning-based image restoration model implemented in AIVIA. To achieve this, we systematically evaluated the effects of training image quality and quantity on the accuracy of image restoration. By testing various combinations of image pairs captured with short-and long-exposure settings, we aimed to identify the optimal training conditions that balanced improving imaging accuracy with minimizing photobleaching and phototoxicity. The long-exposure images (2000 ms) provided high-quality confocal images, while the short-exposure images reduced photobleaching and phototoxicity. This optimization process was crucial for determining the most suitable conditions for subsequent 4D live-cell imaging analysis. To examine the effects of training image quality, we prepared four patterns of training image sets with different short exposure times (Figure 1). We also examined the effect of the number of training image sets on the model’s image restoration accuracy by training several models with different numbers of training image sets. Specifically, we trained 24 models with a combination of four short exposure times (20, 50, 100, and 100 ms) and 6 datasets consisting of different numbers of training image sets (N = 2, 6, 12, 25, 50, and 100) to verify the image restoration accuracy.

**Figure 1.**
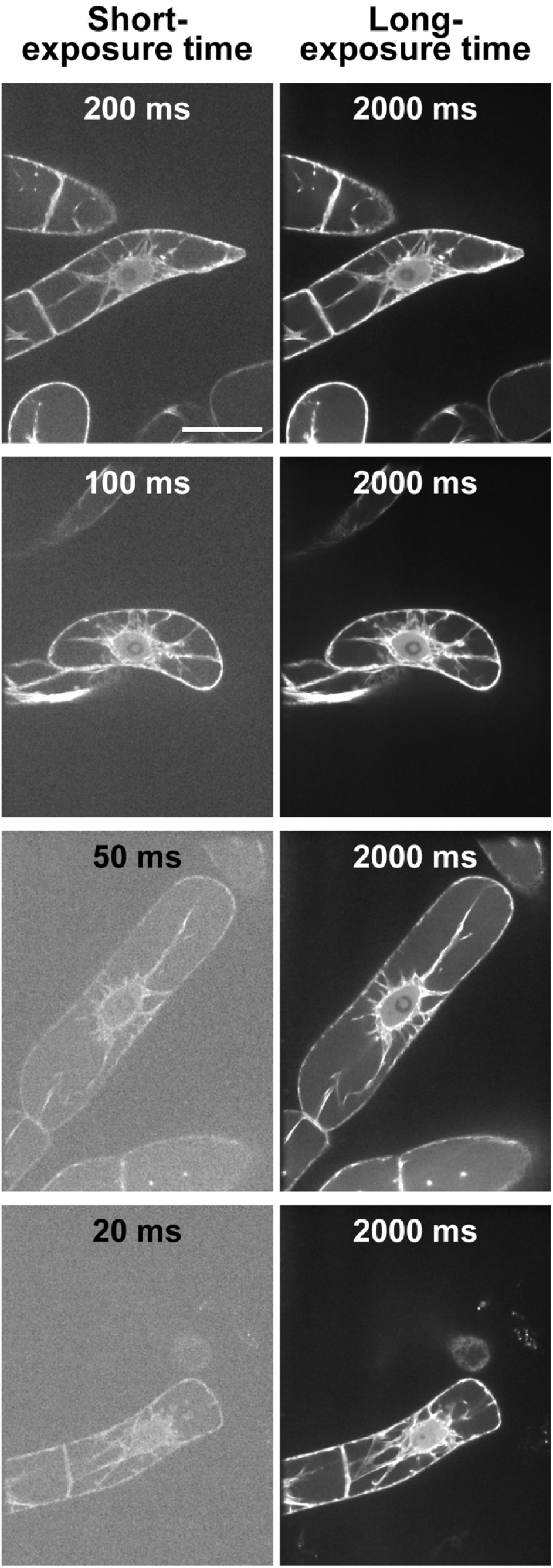
Training image sets for image restoration models. A training image set consists of a low-quality image taken with a range of short exposure times and a high-quality image taken with a long exposure time. There are four combination patterns of training image sets: 200 and 2000 ms images, 100 and 2000 ms images, 50 and 2000 ms images, and 20 and 2000 ms images. Scale bar = 40 µm.

We then evaluated the image restoration accuracy of the 24 models qualitatively and quantitatively by examining the restoration results of the test images (Figures 2, 3). Qualitatively, the image restoration accuracy tended to improve in proportion to the quality and number of training images (Figure 2). In other words, the detected signals were clearer (relative to the background noise) after image restoration using training images with longer exposure times and models with larger numbers of training image sets. For the quantitative evaluation, Spearman’s rank correlation coefficient (SRCC) was calculated to quantify the degree of improvement in similarity to the *ground truth* image (long-exposure image) after image restoration by the model (Figure 3). The results of the quantitative evaluation were consistent with the results of the qualitative evaluation and showed an improvement in restoration accuracy that was almost proportional to the quality and number of training image sets. Quantitative evaluation by SRCC showed two points: (i) SRCC was lower for relatively low-quality training image sets (20 or 50 ms) than for raw input images when the number of training image sets was very small (N = 2) (Figure 3a, b). This tendency was not observed when the training image sets were of relatively high quality (100 or 200 ms) (Figure 3c, d). (ii) SRCC was comparable between images restored using relatively low-quality images with the short exposure time of 50 ms and images restored using higher quality input images, if the number of training image sets was sufficient (N = 100) (Figure 3b–d). The mean and standard deviation (SD) of SRCC for each exposure time (N = 100) were 0.812 ± 0.21 (20 ms), 0.914 ± 0.07 (50 ms), 0.930 ± 0.05 (100 ms), and 0.962 ± 0.03 (200 ms) (Figure 3). This suggests that image restoration using a model trained with a sufficient number of training image sets can produce images with a quality similar to that of the image taken with a long exposure time (2000 ms), even for a low-quality image taken with a short exposure time of 50 ms. On the basis of the above evaluation results, we used the model trained with the combination of “2000 ms long-exposure image, 50 ms short-exposure image, and N = 100 training image set” for image restoration during the analysis of 4D live-cell images of tobacco BY-2 cells, as described later.

**Figure 2.**
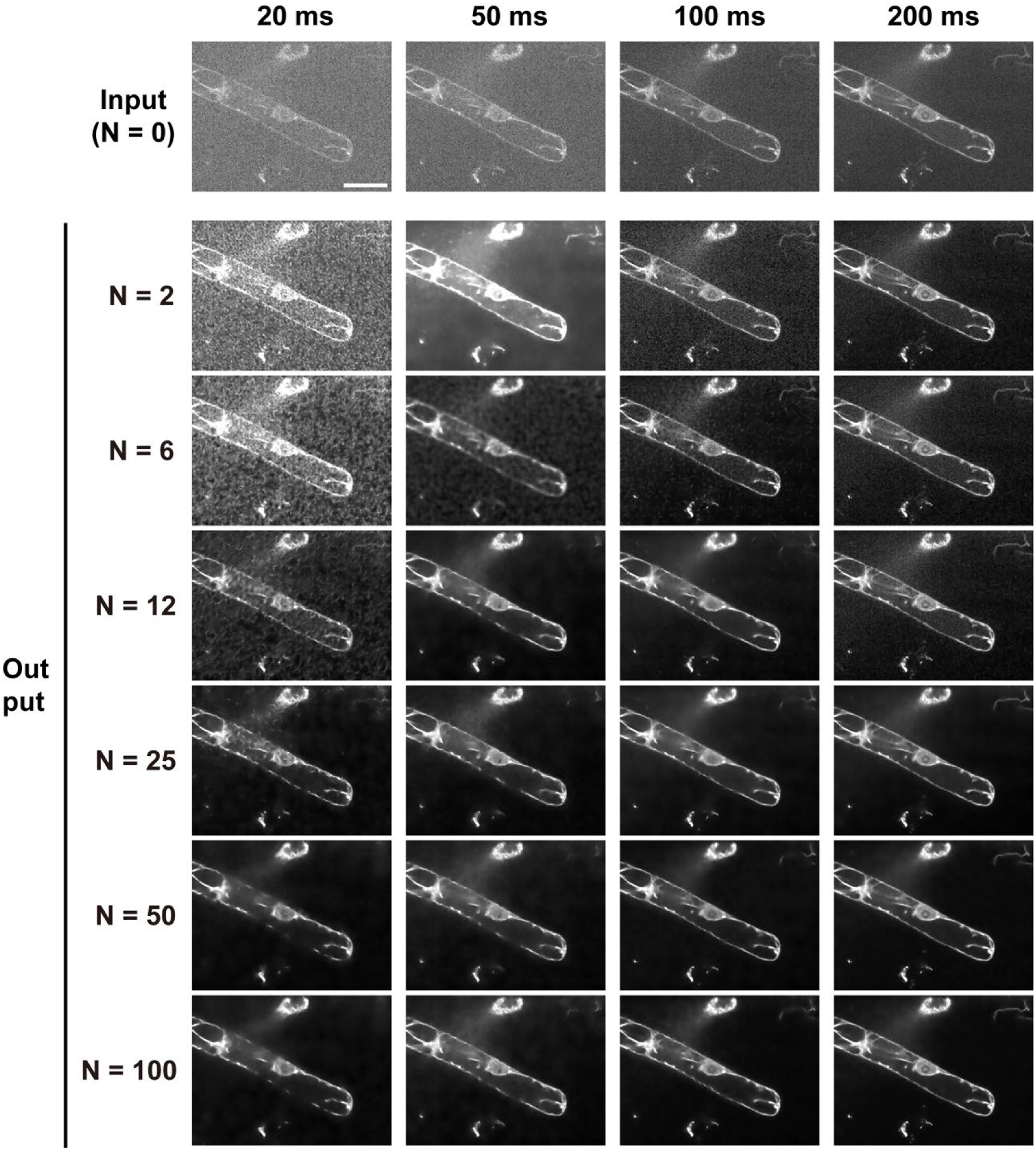
Effect of the quality and number of training image sets on image restoration. “Input” is images of tobacco BY-2 cells in which actin microfilaments are fluorescently labeled with Lifeact-RFP and taken with the following pattern of exposure times: 20, 50, 100, and 200 ms. “Output” is the result of image restoration performed on images taken with the respective exposure times. N indicates the number of training image sets used to train the image restoration model. Scale bar = 40 µm.

**Figure 3.**
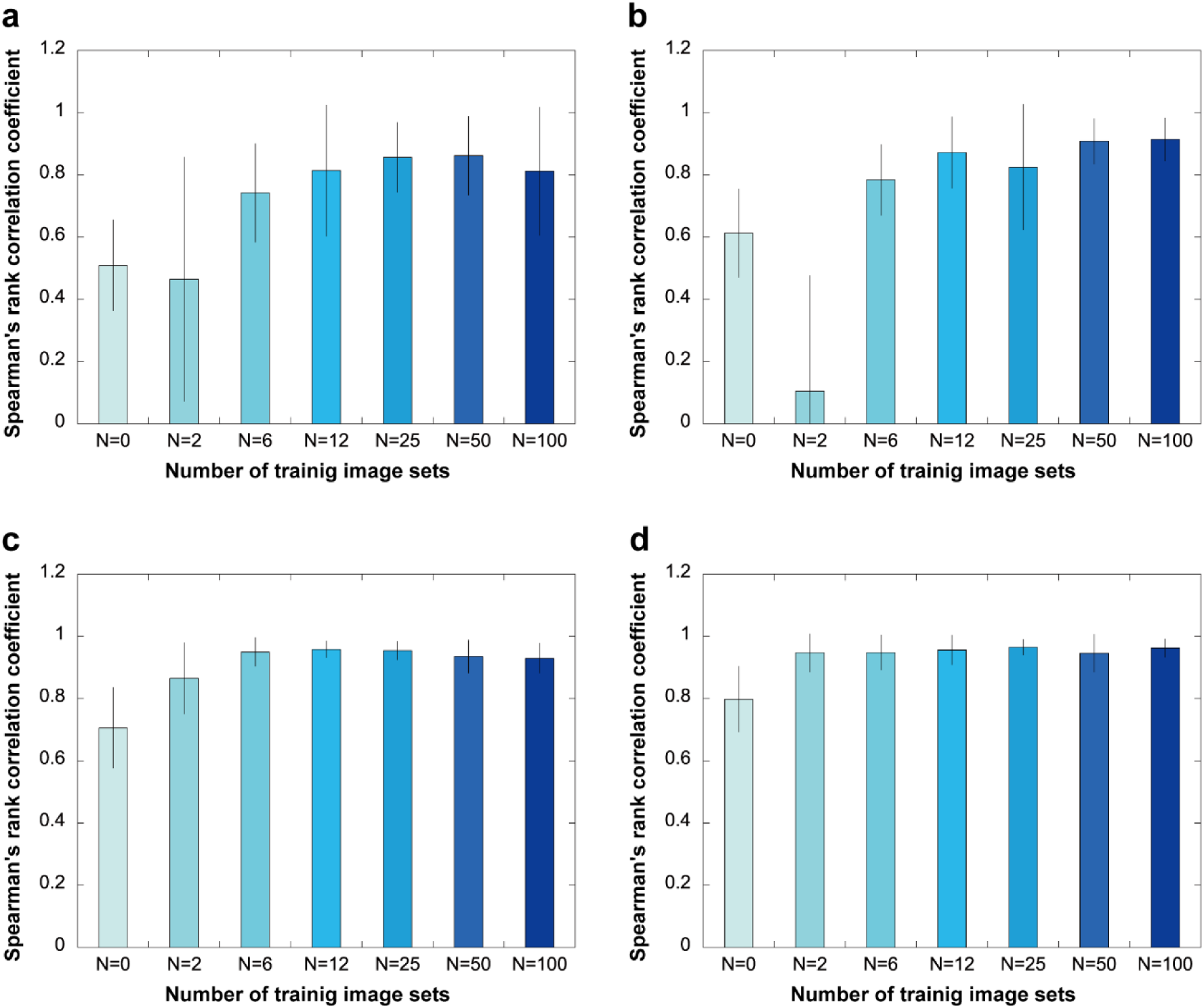
Quantitative evaluation of the effect of the number of training image sets and input image quality on image restoration accuracy. The results shown are from the quantitative evaluation of image restoration performed on Lifeact-RFP fluorescence images. N indicates the number of training image sets. The input images (N = 0) are taken with four patterns of exposure times: (a) 20, (b) 50, (c) 100, and (d) 200 ms. The input image and the images restored with each model (N = 2, 6, 12, 25, 50, and 100) were compared with the *ground truth* image (taken with a long exposure time of 2000 ms), and their similarity was quantified using Spearman’s rank correlation coefficient (SRCC). The vertical axis represents SRCC, with SRCC = 1 indicating high image restoration accuracy. Error bars indicate standard deviation (*n* = 500 for each bar).

To evaluate the utility of the image restoration technique for bioimaging of plant cells other than tobacco BY-2 cells, we extended the validation of image restoration to fluorescence live-cell imaging of zygotes of a model plant, *A. thaliana* (Supplemental Figure 2) (Kimata et al., 2016; Hiromoto et al. 2023). The models were trained using pairs of low-and high-quality images or medium-and high-quality images of *A. thaliana* zygotes. The quality of each image was determined by the excitation power of the applied laser (detailed acquisition conditions are described in Materials and Methods). The model optimized for *A. thaliana* zygotes improved image restoration accuracy both qualitatively and quantitatively (Supplementary Figure 2). The mean ± SD of the SRCCs were 0.12 ± 0.02 for input-low, 0.68 ± 0.07 for output-low, 0.20 ± 0.03 for input-medium, 0.70 ± 0.06 for output-low (Supplementary Figure 2b). For both the low-and medium-quality images, there was a significant improvement in the quality of the output image compared with that of the input image. In other words, the output image was more similar to the *ground truth* high-quality image than the input image (Supplementary Figure 2).

### Image restoration improves the quality of 3D reconstructed images of AFs in tobacco BY-2 cells acquired by low-damage imaging with short exposure times

We then applied the image restoration model to the observation of AF dynamics in the initial phase of cell plate formation (early telophase) and performed 3D reconstruction of Z-stack images at selected time points (Figure 4). In early telophase, Lifeact-RFP fluorescence was enhanced in the output image (Supplemental Video 1) compared with the input image taken with a short exposure time (50 ms) (Supplemental Video 2). This enhancement confirmed the presence of Lifeact-RFP in the inner region where cell plate formation occurs (Figure 4, fifth row in early telophase). The signal-to-noise ratio was improved in the output image compared with the input image, making it suitable for 3D structural analysis. Because enhancing the fluorescence signal by increasing the exposure time often leads to photobleaching and phototoxicity (Supplemental Figure 1), enhancing the fluorescence signal in short-exposure images by image restoration can be a valuable tool in low-damage live-cell observations. In late telophase, the improvement in image quality by image restoration was modest compared with early telophase. The output image showed a slight noise reduction compared with the input image, and it showed a Lifeact-RFP signal localized at the outer edge of the ring-shaped phragmoplast, like the input image (Figure 4, fourth and fifth row in late telophase).

**Figure 4.**
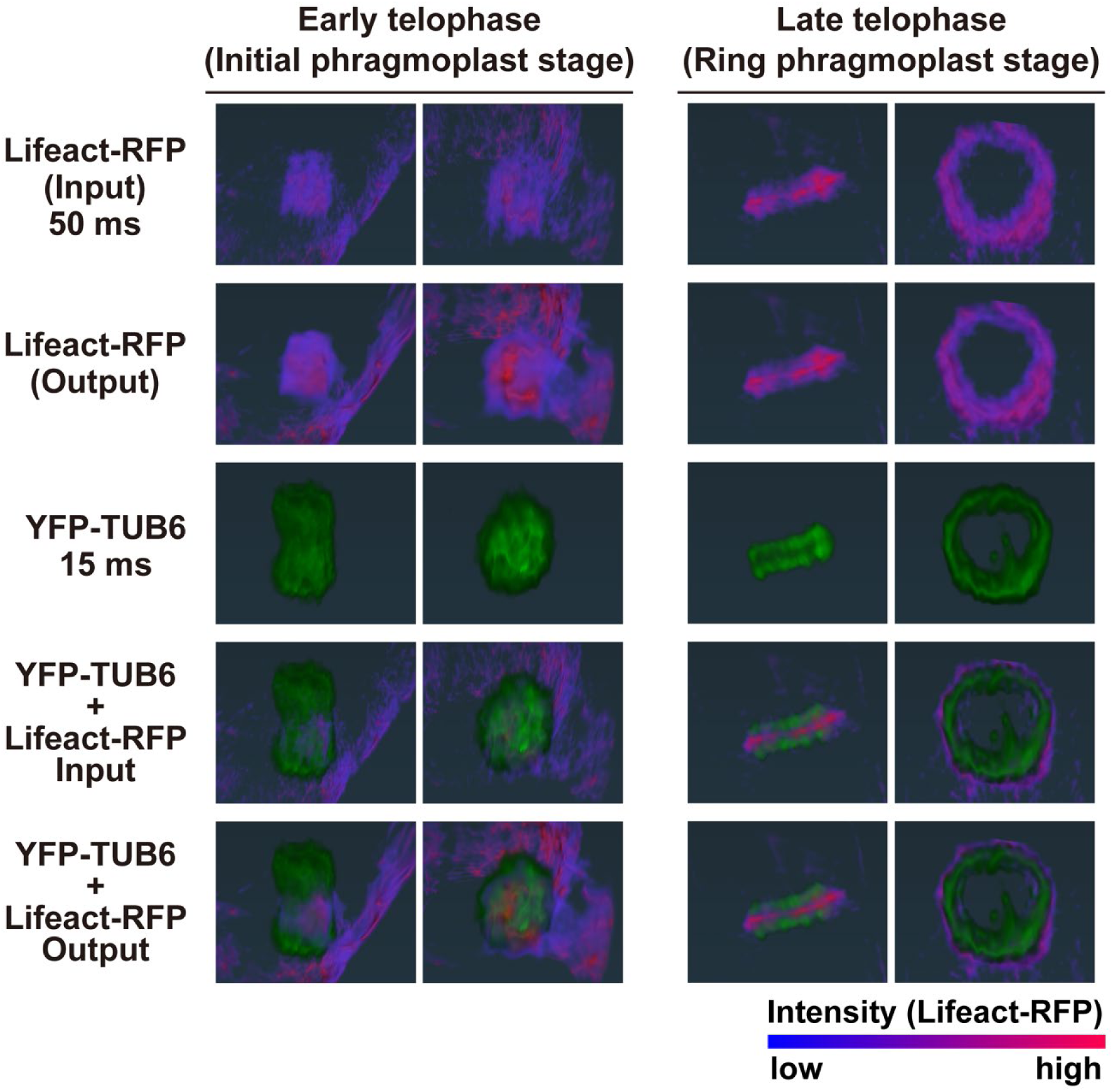
Effect of image restoration on 4D imaging of tobacco BY-2 cells. Three-dimensional reconstruction of Z-stack images acquired during early telophase and late telophase at the corresponding timepoints. For each phase, the right column shows a vertical view of the cell plate formation plane of the object in the left column. Microtubules labeled with YFP-TUB6 and actin microfilaments labeled with Lifeact-RFP are shown in pseudo colors, with the increase in fluorescence intensity from image restoration indicated by the color map of Lifeact-RFP pseudo colors.

### Lifeact-RFP labeled AFs predominately localize near the cell plate during initial stage of formation

To more precisely investigate the localization and dynamics of Lifeact-RFP labeled AFs during the initial phase of cell plate formation, as indicated by 3D analysis, we examined temporal changes in optical sections through the midplane of the mitotic apparatus (Figure 5a). Owing to the noise reduction and AF-signal enhancement by image restoration, time-lapse images taken at intervals of 30 s clearly showed the localization of Lifeact-RFP-labeled AFs at the site of cell plate formation (Figure 5a, arrowheads at 0 s in the third and fourth rows). Lifeact-RFP signals near daughter nuclei were also detected as the cell plate expanded (Figure 5a, 30–90 s in the third and fourth rows). These time-lapse images are composed of optical sections of Z-stack images acquired at each time point.

**Figure 5.**
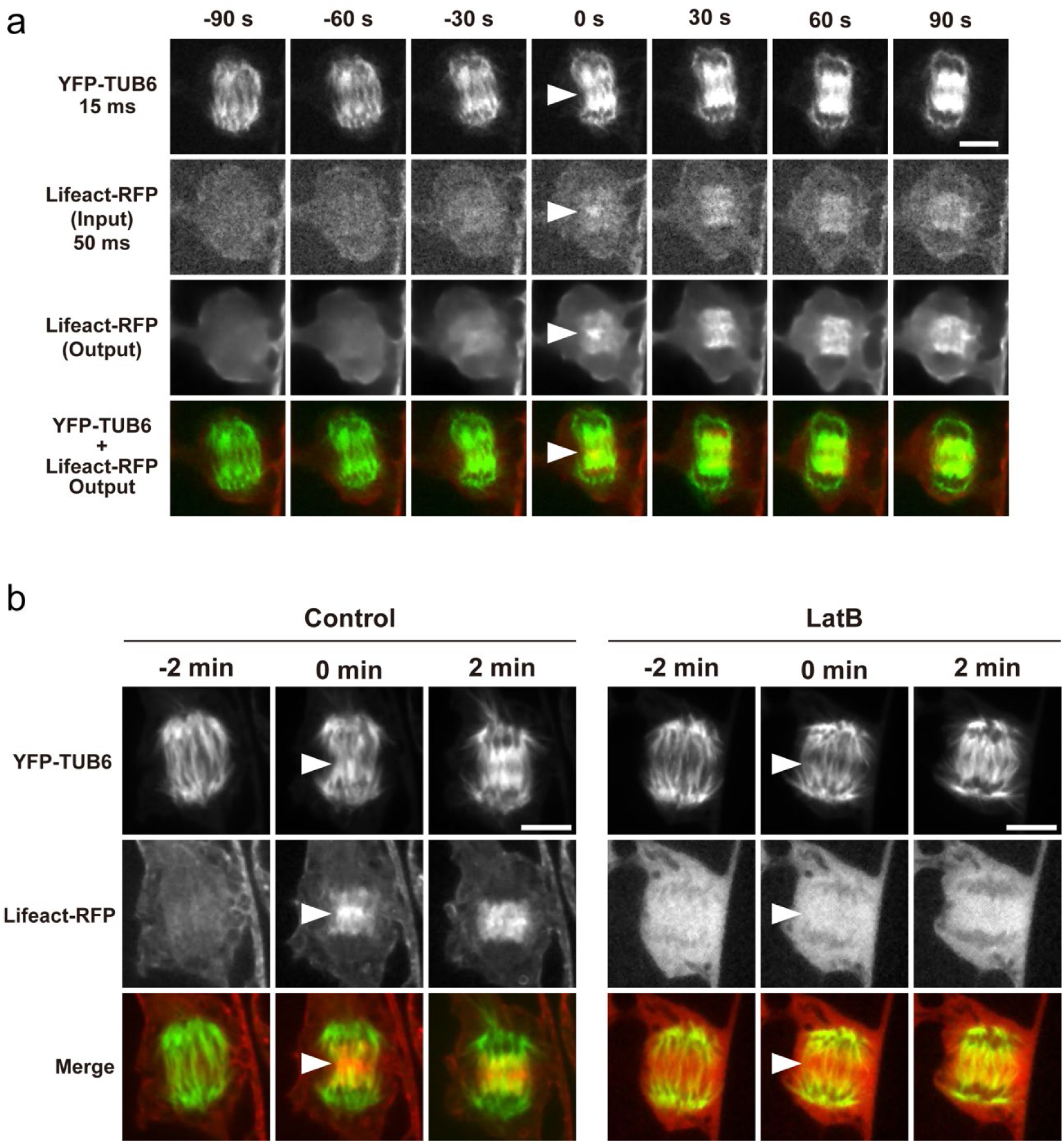
Application of image restoration to the observation of tobacco BY-2 cells with fluorescently labeled actin filaments (Lifeact-RFP) and microtubules (YFP-TUB6). (a) Time-lapse observation using image restoration. (b) Images taken with a long exposure time. Cells were untreated (Control) or treated with 4 µM Latrunculin B for 1−2 h (LatB) to inhibit actin polymerization. Each time-lapse image is shown with a representative single optical section through the midplane of the division apparatus of 41 (a), 8 (b, Control), or 7 (b, LatB) cells. The time point at the onset of cell plate formation is indicated as 0 min. Arrowheads indicate the initiation region of cell plate formation. Scale bars = 10 µm.

Careful examination of the structures of interest found in the restored images is essential because image restoration may produce artifacts from the addition of extra signals or removal of signals from the images. We then checked long-exposure images taken at the same phase (or initial phase of cell plate formation) (Figure 5b). It should be noted that long-term time-lapse imaging with long exposure times results in cell cycle arrest owing to phototoxicity and fading of the fluorescence signal owing to photobleaching (Supplemental Figure 1). Therefore, we performed long-exposure imaging at 3−5 timepoints in the phase of interest, which were pre-determined by observations using image restoration. Because the Z-stack images were acquired with long exposure times, the shortest interval was 2 min. These time-lapse images provided optical sections through the midplane of the mitotic apparatus (Figure 5b). AF localization at the site of cell plate formation was directly observed in the long-exposure images, confirming that AF localization inferred through image restoration corresponded to actual structures (Figure 5a, b, Control).

We also tested the inhibition of actin polymerization by LatB treatment to verify whether the signal of Lifeact-RFP labeled AFs detected at the site of cell plate formation was derived from AFs (Figure 5b, LatB). After 1−2 h of LatB treatment, the Lifeact-RFP signal was diffused around the mitotic apparatus, indicating that actin polymerization was inhibited (Figure 5b, LatB). Comparison of untreated (Figure 5b, Control) and LatB-treated (Figure 5b, LatB) groups showed that the signal of Lifeact-RFP labeled AFs at the site of cell plate formation was indeed derived from AFs.

### Two different probes of AFs, Lifeact-RFP and RFP-ABD2, localize differently during initial stage of cell plate formation

To determine whether AFs labeled with other fluorescent probes exhibit the same localization pattern and dynamics as AFs labeled with Lifeact-RFP, tobacco BY-2 cells were fluorescently labeled with RFP-ABD2 and analyzed. The sequence of the analysis was identical to that for Lifeact-RFP labeled AFs, consisting of time-lapse imaging with image restoration, validation using images taken with long exposure times, and actin polymerization inhibition assays (Figure 6).

**Figure 6.**
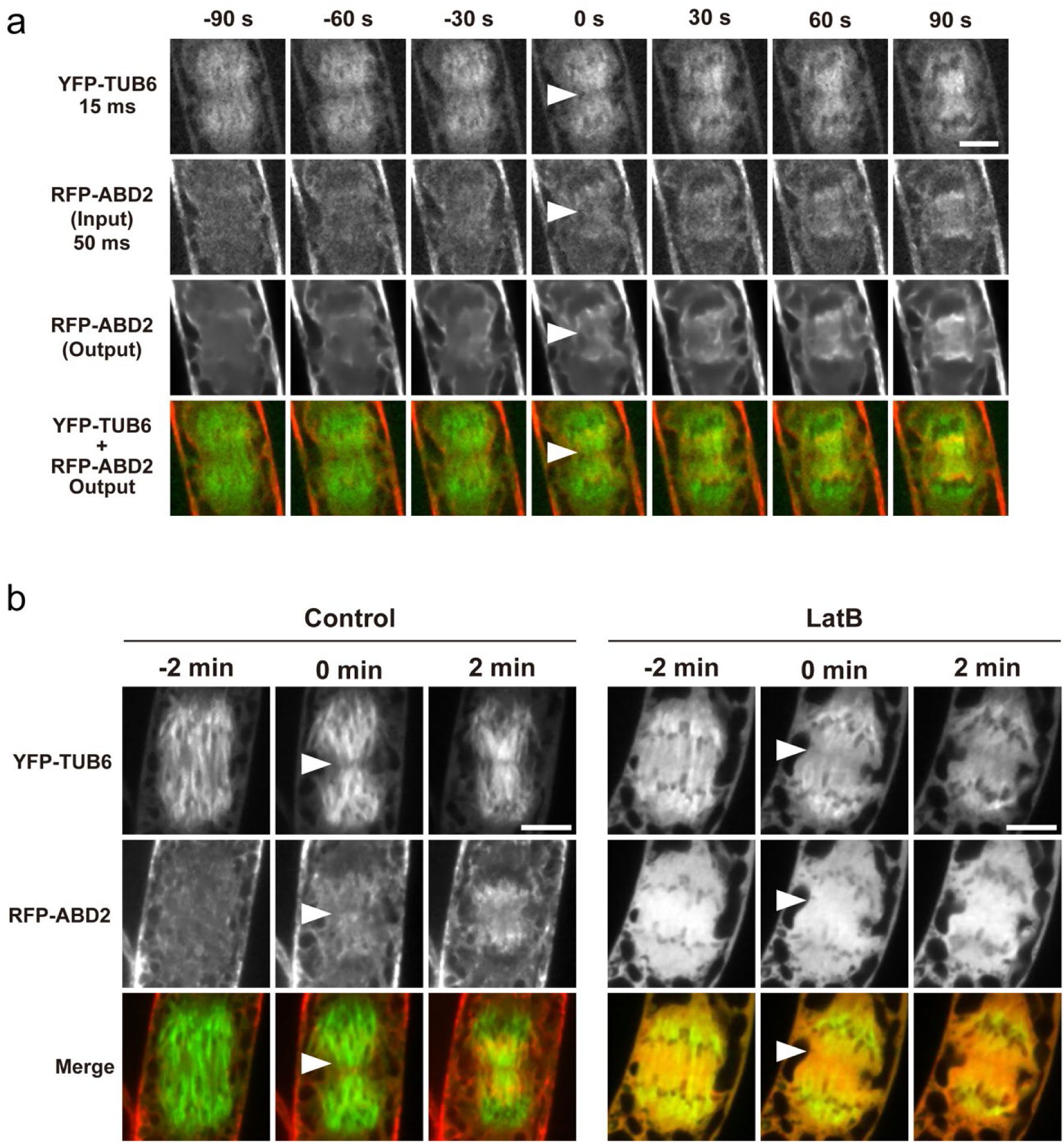
Application of image restoration to the observation of tobacco BY-2 cells with fluorescently labeled actin filaments (RFP-ABD2) and microtubules (YFP-TUB6). (a) Time-lapse observation using image restoration. (b) Images taken with a long exposure time. Cells were untreated (Control) or treated with 4 μM Latrunculin B for 1−2 h (LatB) to inhibit actin polymerization. Each time-lapse image is shown with a representative single optical section through the midplane of the division apparatus of 36 (a) or 14 (b, Control), or 6 (b, LatB) cells. The time point at the onset of cell plate formation is indicated as 0 min. Arrowheads indicate the initiation region of cell plate formation. Scale bars = 10 µm.

Image restoration of time-lapse images of RFP-ABD2 labeled AFs was performed using the model already established in the previous section (i.e., the model trained on sets of Lifeact-RFP images acquired at 2000 and 50 ms). As shown in Figure 6a, the model trained on Lifeact-RFP images successfully improved the quality of time-lapse images of RFP-ABD2 labeled AFs (Figure 6a, second and third rows). Time-lapse images with image restoration showed that RFP-ABD2 labeled AFs rarely localized near the cell plate in the early stage of plate formation (Figure 6a, 0 s in third and fourth rows) but localized near the daughter nuclei as the cell plate expanded (Figure 6a, 30−90 sec in the third and fourth rows). This observation is consistent with previous reports (Higaki et al., 2008; Maeda et al., 2020).

Time-lapse images of RFP-ABD2 labeled AFs acquired with long exposure times supported the localization pattern detected by image restoration, as in the case of Lifeact-RFP (Figure 6b, Control). In addition, inhibition of actin polymerization by LatB treatment disturbed the localization pattern of RFP-ABD2 labeled AFs (Figure 6b). Taken together, our results showed that image restoration revealed accurate dynamics of AFs, even in the case of RFP-ABD2 labeled AFs, shedding light on the differences in the dynamics of AFs labeled with Lifeact-RFP and RFP-ABD2 (Figure 7).

**Figure 7.**
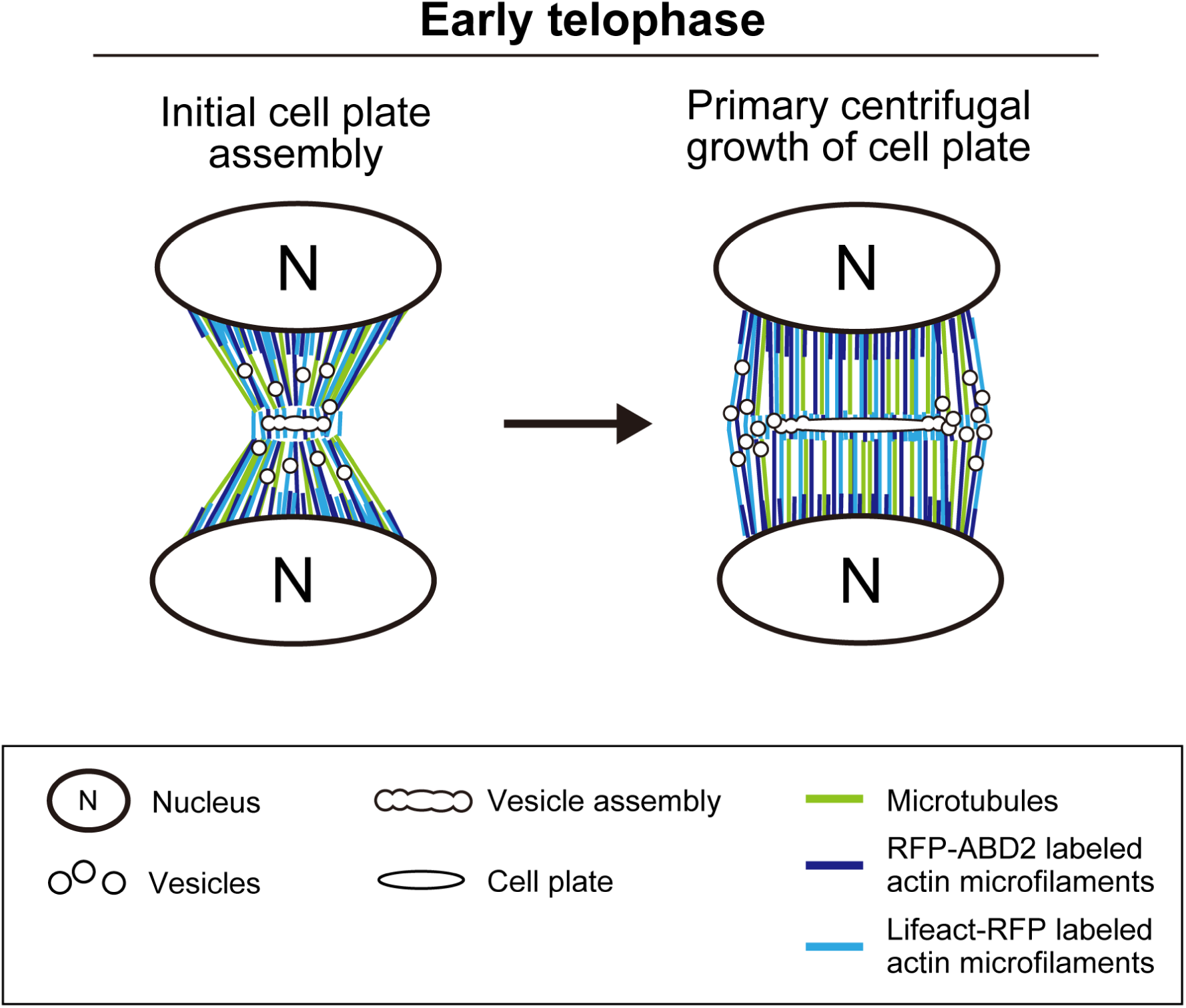
Schematic diagram of actin microfilament dynamics during early telophase. Lifeact-RFP and RFP-ABD2 labeled actin microfilaments localized around the cell plate initiation region and daughter nuclei during initial cell plate assembly, to different levels. The vesicle assembly region was dominated by Lifeact-RFP labeled actin microfilaments, with few or absent RFP-ABD2 labeled actin filaments. During primary centrifugal growth of the cell plate or phragmoplast expansion, Lifeact-RFP labeled actin microfilaments were highly co-localized with microtubules of the phragmoplast and localized around the forming cell plate, while RFP-ABD2 labeled actin microfilaments were highly localized near the daughter nuclei.

## Discussion

### Image restoration is an effective tool for reducing the negative effects of laser exposure and improving spatiotemporal resolution in live-cell imaging of tobacco BY-2 cells

In this study, we applied deep learning-based image restoration models to improve the quality of fluorescent images of AFs in tobacco BY-2 cells during cell division. Deep learning has several advantages for image restoration in live-cell imaging. Deep learning algorithms can effectively remove various features degrading images, such as noise, blur, and artifacts, resulting in clearer and more accurate images (Belthangady and Royer, 2019). Additionally, deep learning-based image restoration can be easily applied to different imaging modalities and datasets, making them versatile and adaptable to various research needs (Jin et al., 2020; Huang et al., 2023; Ma et al., 2024). In fact, deep learning-based image restoration has recently been used to improve the spatiotemporal resolution of fluorescence live-cell imaging (Weigert et al., 2018; Wang et al., 2019; Laine et al., 2021). Because this technology enhances the weak signals buried in noise, laser exposure can be minimized during imaging (Chen et al., 2020; Laine et al., 2021; Boothe et al., 2023; Ma et al., 2024). However, there are also some disadvantages to using deep learning for image restoration. Deep learning algorithms heavily rely on training data, and the quality and diversity of the training dataset can significantly impact the performance of the restoration model (Belthangady and Royer, 2019). It is often difficult to experimentally acquire large numbers of appropriate training image sets (low-and high-quality image pairs), which is also an obstacle to the practical application of deep learning-based fluorescence image restoration. Moreover, deep learning models applied to extremely noisy images may introduce artifacts or deceiving results, which affects the accuracy of quantitative measurements and tracking results (Belthangady and Royer, 2019). Hence, we qualitatively and quantitatively evaluated the accuracy of the image restoration method used (Figures 2, 3), as described below.

Our qualitative and quantitative evaluations revealed that the quality and number of training image sets are associated with image restoration accuracy (Figures 2, 3). It is interesting to note that using a model trained on 100 image sets to restore an image taken with an exposure time of 50 ms produced an output image that was similar to the *ground truth* image acquired with an exposure time of 2000 ms (Figure 3b). Features of AFs labeled with Lifeact-RFP or RFP- ABD2, as indicated by image restoration, also existed in images taken with a long exposure time (Figures 5, 6). Thus, our image restoration model enabled high-resolution live-cell imaging with reduced exposure times, from 2000 ms to 50 ms, minimizing damage to tobacco BY-2 cells. Moreover, the image restoration model was trained on a relatively small number of image sets using the pre-trained model in AIVIA, demonstrating the potential to apply image restoration to the observation of objects for which a large number of training image sets was difficult to acquire. It should be noted that the accuracy of restoring short-exposure images decreased significantly when the number of training image sets was extremely small (Figure 3a, b; number of training image sets = 2, exposure time of input images = 20 and 50 ms), which may be related to artifacts arising from noisy features in the training images, as described in previous studies (Belthangady and Royer, 2019).

### Limitations and potential applications of image restoration in this study

This study expands the potential use of image restoration models in biological research, particularly in live-cell imaging of plants. However, it should be noted that acquiring a suitable image set for training the image restoration model is a major limiting factor. That is, when acquiring high-and low-quality image sets, if the positions of the objects of interest in the images do not match, the accuracy of the model after training will significantly degrade. This problem is critical when acquiring training image sets of models focusing on intracellular structures that rapidly change their shape.

Future applications of image restoration include exploratory observations of unknown intracellular phenomena. Although long-term live-cell imaging with short acquisition intervals are effective for detecting the dynamics of novel and interest intracellular structures, they are difficult to conduct in practice owing to the negative effects of prolonged observation, namely phototoxicity and photobleaching (Magidson and Khodjakov, 2013; Skylaki et al., 2016; Ojha and Ojha, 2021). This study showed that image restoration enabled live-cell imaging with reduced phototoxicity (e.g., cell cycle arrest) and photobleaching (Supplementary Figure 1), paving the way for long-term live-cell fluorescence imaging with reduced cellular damage. This positions our image restoration approach alongside optical approaches that minimize laser exposure, such as light sheet microscopy and lattice light sheet microscopy (Daetwyler and Fiolka, 2023; Sinclair et al., 2024). High-resolution 4D live-cell imaging with reduced cellular damage using image restoration also enables the detection of various biological events for insights into cellular workings. Our image restoration approach was applicable to live-cell imaging of not only tobacco BY-2 cells but also *A. thaliana* zygotes (Supplementary Figure 2), suggesting that it could be applied to study a wide range of plant species and cell types, if suitable training images are available.

### Differences in the dynamics of AFs labeled with different probes in the initial phase of cell plate formation

Analyzing 4D images of tobacco BY-2 cells using our image restoration technique revealed that the dynamics of AFs in the initial phase of cell plate formation differed depending on the attached probe. Specifically, the vesicle assembly region was dominated by Lifeact-RFP labeled AFs, with little or absent RFP-ABD2 labeled AFs, which were predominately localized near the daughter nuclei (Figures 5−7). Previous studies reported that AFs are involved in the initial phase of cell plate formation (Higaki et al., 2008; Maeda et al., 2020; Maeda and Higaki, 2021). The different dynamics of Lifeact-RFP labeled AFs and RFP-ABD2 labeled AFs suggest that specific types of AFs are involved in vesicle trafficking during cell plate formation (Figures 5−7).

The difference in dynamics between Lifeact-RFP-labeled AFs and RFP-ABD2-labeled AFs may be understood by considering the respective properties of Lifeact and ABD2. ABD2 fused with fluorescent proteins has been used to visualize the dynamic organization of AFs *in vivo*, providing detailed structural insights (Sheahan et al., 2004; Wang et al., 2008; Higaki et al., 2008; Maeda et al., 2020). Lifeact, a peptide consisting of the first 17 amino acids of yeast protein Abp140, has been used to clearly visualize filamentous AFs in eukaryotic cells and tissues. (Riedl et al., 2008). Previous studies have shown that Lifeact is an excellent probe for studying AF organization and dynamics in plants because, when expressed to the correct level, it labels AF populations such as thin F-actin, which are undetectable with ABD2 as the AF marker (van der Honing et al., 2011). However, certain native AFs of plant cells may remain unaffected by Lifeact because a number of AtACT2-containing filaments in *Nicotiana benthamiana* leaf epidermal cells do not interact with Lifeact (Kijima et al., 2018). Moreover, different actin isoforms can build their own filament arrays depending on the cell and tissue type, and selective visualization of each filament array is possible depending on the type of actin-binding protein, as proposed by Kijima et al. (2018). Considering the above studies, it is possible that AFs specifically labeled with Lifeact-RFP, rather than RFP-ABD2, regulate vesicle arrays *in situ* in the initial phase of cell plate formation. Actin polymerization inhibition tests with LatB showed that the midzone contraction of the anaphase spindle is inhibited in the initial stage of cell plate formation (Maeda et al., 2020), which may be attributed to the absence of Lifeact-RFP labeled AFs around the midzone and the failure of MT guidance. Future studies focusing on AFs specifically labeled with Lifeact-RFP will provide a deeper understanding of the role of AFs in cell plate formation.

## Author Contributions

T.H. conceived and designed the experiments. S.K., T.K., and T.H. performed 4D imaging analysis using the image restoration model. Y.H. and M.U. acquired the image of *A. thaliana* zygotes. S.K. and T.H. wrote the manuscript, and all authors participated in the discussion and review of the manuscript.

## Acknowledgments

We thank Ms. Hitomi Okada (Kumamoto University) and Ms. Remi Kawakami (Kumamoto University) for their support in cell culture maintenance. We thank Edanz (https://jp.edanz.com/ac) for editing a draft of this manuscript.

## Data Availability Statement

All relevant data are available from the authors on request.

---

**Supplemental Figure 1.**
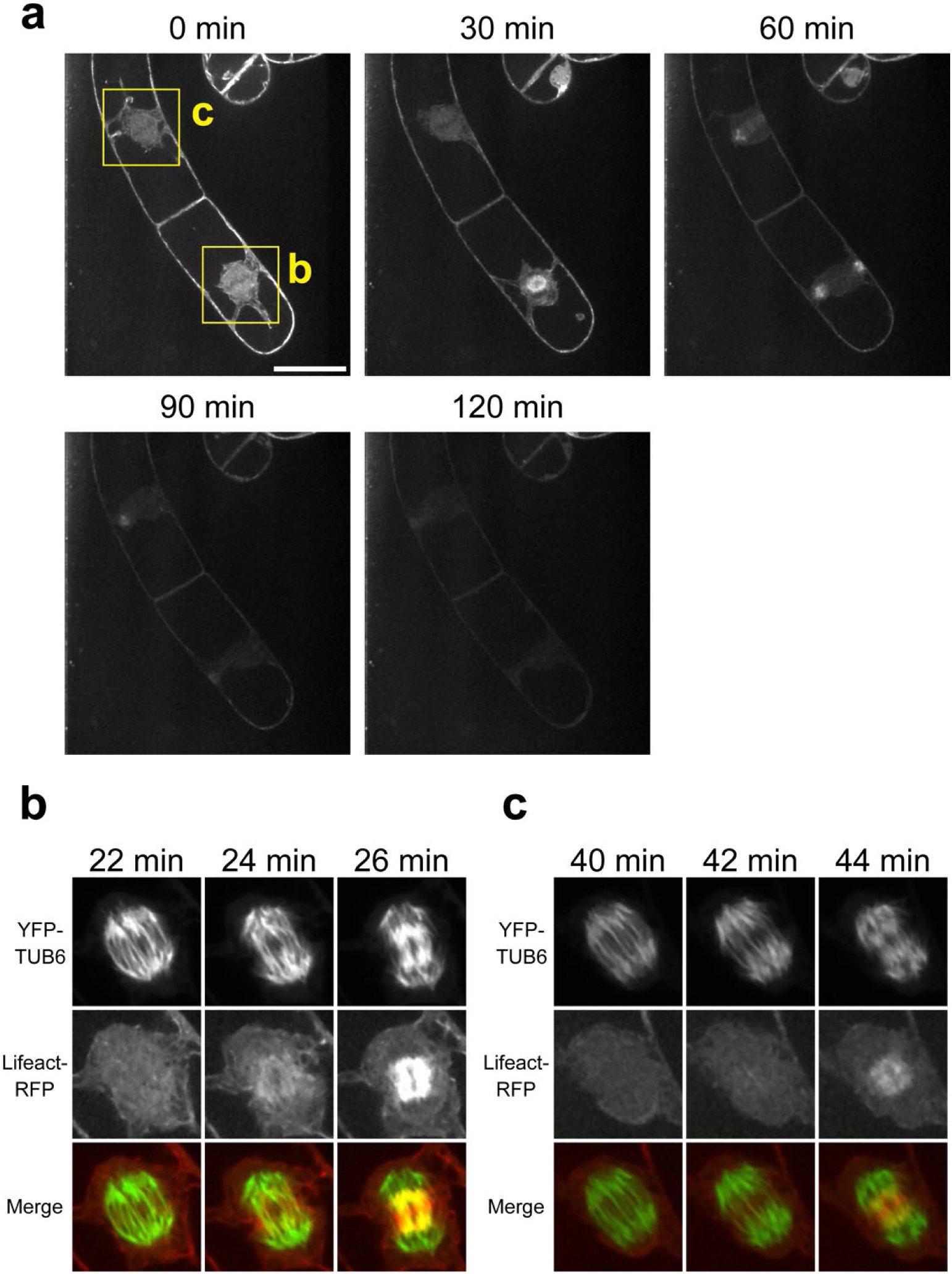
Photobleaching during live-cell fluorescence imaging. (a) Time-lapse images of tobacco BY-2 cells with fluorescently labeled microtubules (YFP-TUB6) and actin microfilaments (Lifeact-RFP). (b) and (c) are enlarged images of the mitotic apparatus surrounded by the yellow squares in (a). The acquisition time interval was 2 min, and 40 optical sections (Z-interval of 1 µm) were obtained at each time point. Each time-lapse image is shown with a representative single optical section through the midplane of the division apparatus of four cells. The time point of the start of acquisition is indicated as t = 0. (b) Scale bar = 40 µm.

**Supplemental Figure 2.**
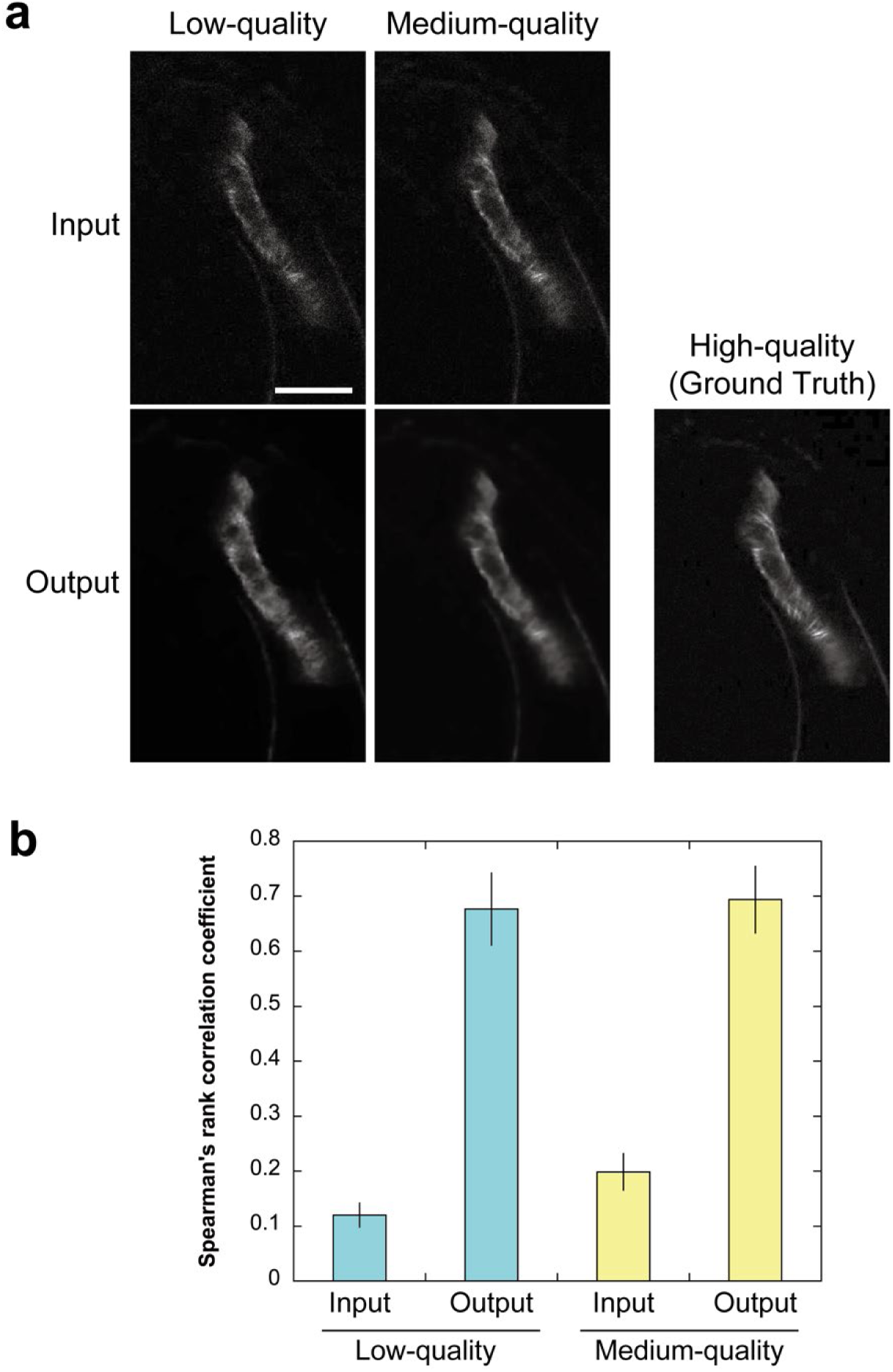
Application of image restoration to the imaging of *A. thaliana* zygotes. (a) Representative images of *A. thaliana* zygote with fluorescently labeled microtubules (Clover-TUA6) before and after image quality restoration. Results for low-(left) and medium-quality (center) images are shown with “Input” (upper) and “Output” (lower) images. The corresponding high-quality image is shown in the lower right. (b) Results of the quantitative evaluation of image restoration accuracy in which “Input” and “Output” images were compared with the *ground truth* image, and their similarity was quantified using Spearman’s rank correlation coefficient (SRCC). Error bars indicate standard deviation (*n* = 23 for each bar). Scale bar = 20 µm.

